# Choice of annotation data significantly impacts automated reconstruction of genome-scale metabolic networks

**DOI:** 10.64898/2026.11.05.764556

**Authors:** B. Adam Bates, Jason A. Papin

## Abstract

Genome-scale metabolic models are valuable tools for studying metabolism in many applications, such as metabolic engineering or understanding human disease. However, curated models are available for only a limited number of organisms, and generating high-quality models for additional organisms remains a challenge. To help address this challenge, several software tools have been developed that can produce draft models in an automated fashion, but the quality of these draft models lags behind manually curated ones. Thus, there is still a need to improve these tools. Previous work has focused primarily on developing better methods for each step in the reconstruction process, such as gene annotation and gap-filling. Here, we focus on the data that underlies these methods, such as known connections between protein sequence and function. We find that changing the source of reconstruction data has a substantial impact on the contents and predictions of generated models. In fact, the source of data can have a bigger impact than the choice of reconstruction method. Additionally, we show that the reconstruction process is sensitive to the accuracy of gene annotations in the reconstruction data. Finally, we demonstrate the importance of having functional annotation data from an organism that is closely related to the one for which a model is being built. These observations highlight the importance of reconstruction data and identify principles that should be considered when selecting data for reconstruction tools.

**Author Summary:** Genome-scale metabolic models are helpful tools for studying the chemical reactions that take place inside of cells. Unfortunately, building these models is challenging. Past efforts have produced software tools that can build metabolic models in an automated fashion; however, the quality of models produced by these tools is still lower than manually curated models. Rather than focusing on the algorithms or methods in these tools, we instead explore the importance of the data that these tools rely on, which we call “reconstruction data”. We show that building models with different reconstruction data can have a bigger impact on the quality of the models than using a different method. Additionally, we demonstrate that the quality of generated models is sensitive to both the accuracy of the data and the organisms that are represented in the data. These results emphasize the importance of reconstruction data and indicate that focusing more attention on reconstruction data has the potential to substantially improve the quality of automated network reconstruction.

## Introduction

Genome-scale metabolic models are valuable tools for studying cellular metabolism at a systems level (1,2). They are used in various applications from guiding metabolic engineering to developing mechanistic hypotheses for specific diseases (1,3). For example, metabolic models were recently used to identify sex-specific changes in ileal metabolism for patients with Crohn’s disease (4). In another recent study, metabolic models were used to identify metabolic interactions within the vaginal microbiome that are associated with bacterial vaginosis (5). As metabolic models and their associated methods continue to improve, even more applications are becoming available (6).

In order to facilitate the use of metabolic models in a wider range of applications, significant effort has been put into automating the steps to generate simulation-ready models (Fig 1A) (7,8). Much of these efforts have focused on improving the methods and algorithms for each step of the process (9,10). One important step is predicting the functions of the genes in an organism for which a model is being built. Several methods have been developed for this step, ranging from assigning gene function based on BLAST sequence similarity (11) to more recent, deep-learning-based approaches such as CLEAN (12). After the predicted annotations have been used to generate a draft network, another important step is gap-filling, which ensures that a model can produce biomass on defined media *in silico* (13). There are multiple approaches for this step, including parsimonious flux-based gap-filling (11), likelihood-based gap-filling (14), and deep-learning-based approaches such as CHESHIRE (15). Ultimately, the efforts to improve model generation have led to several automated tools capable of reconstructing metabolic networks from annotated genomes (10,11,16–24).

**Fig 1.**
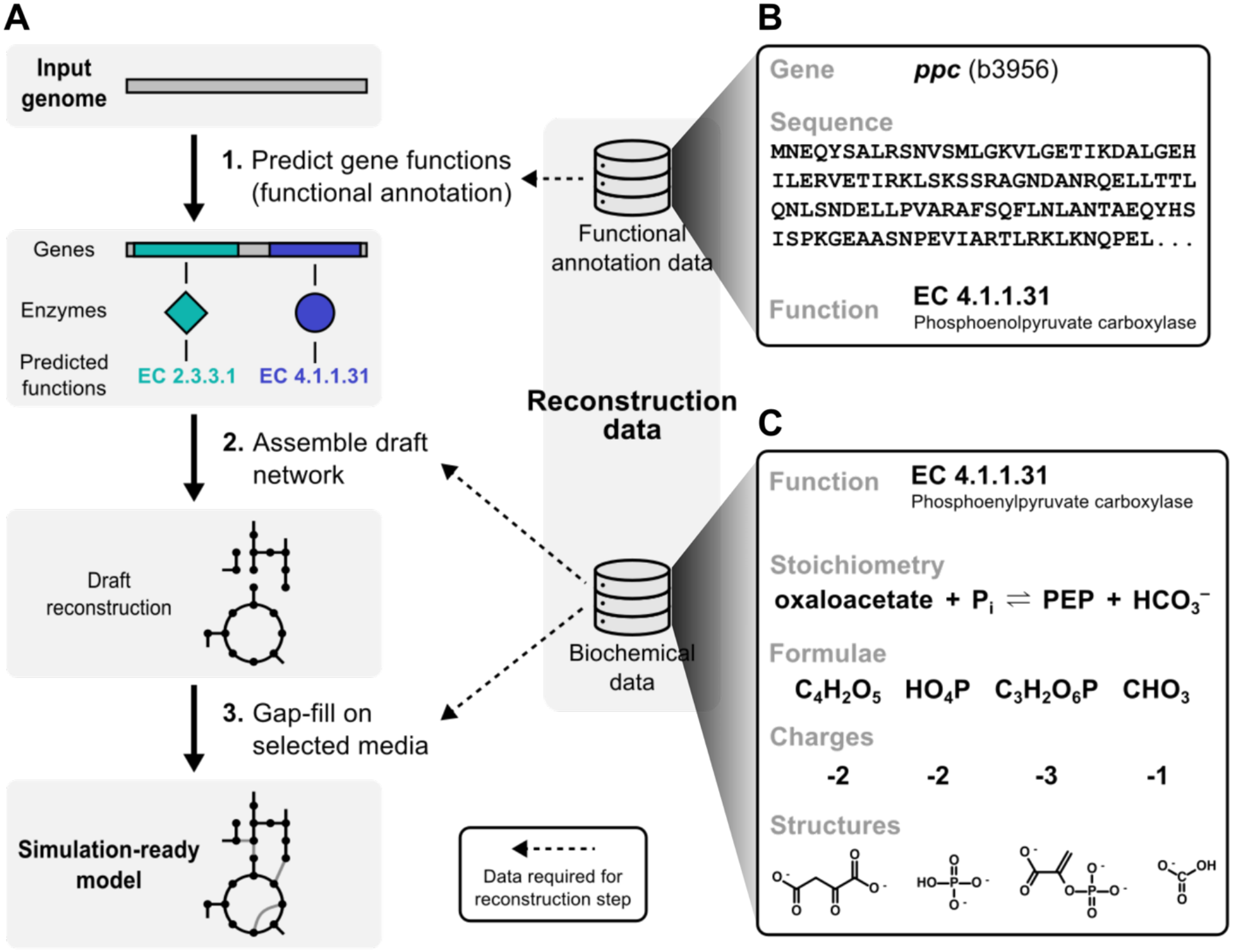
Overview of the process and reconstruction data required to create a metabolic network model. (A) Illustration of the primary steps in automated reconstruction of a genome-scale metabolic network. Many reconstruction tools start from an annotated genome, map the annotated genes to biochemical reactions, create a draft metabolic network reconstruction, and finally perform gap-filling to ensure the model can grow in a specific environment. The reconstruction process requires two main kinds of data: functional annotation data and biochemical data. (B) Example functional annotation data, which link genes to the protein sequence and enzyme function of their gene product. (C) Example biochemical data, which specifies the stoichiometry of biochemical reactions as well as the chemical formulae, charges, and structures of the participating molecules.

One critical component used in the construction of a metabolic model is prior functional annotation data and biochemical data, which we refer to more generally as reconstruction data (Fig 1) (7,25). Here we define “functional annotation data” as known connections between genes and the catalytic function of their gene products (Fig 1B). Additionally, “biochemical data” are data about biochemical reactions and the chemical compounds that are involved (Fig 1C). These data are used throughout the process of reconstructing a metabolic network. For example, methods to predict gene function require functional annotation data, and gap-filling algorithms require biochemical data (25). There are multiple sources for these kinds of genetic and biochemical data such as BiGG (26), KEGG (27), UniProt (28), ModelSEED (29), and MetaCyc (30). While the importance of using high-quality data has been acknowledged (7,16), each automated reconstruction tool typically uses data from only one or two of these sources, and there is little rationale or evidence for the choice of one data source over another. For example, a survey of several published reconstruction tools checked whether each tool used data from an updated database, but no consideration was given to the impact of choosing one database over another (31). Additionally, while different tools have been shown to produce significantly different reaction networks, it is unclear whether such variation in outputs is due to a difference in methods, data, or both (31,32). Thus, the importance of reconstruction data in determining the quality of generated models needs to be established, and principles need to be identified that can guide the selection of these data.

In this work, we explore the importance of reconstruction data and how specific characteristics of the data impact the quality of reconstructed metabolic networks. We show that the source of reconstruction data has a large impact on the contents and predictions of generated models. Since the data from distinct sources differ considerably from each other, we subsequently investigate the effect of certain parameters of the data, such as the accuracy of gene annotations and representation of diverse organisms. We find that both the contents and predictions of models are sensitive to changes in gene annotations. In fact, changing even a single gene annotation in the reconstruction data can greatly alter the contents and predictions of a generated model. We also observe that the reconstruction process is sensitive to what organisms are represented in the reconstruction data. This impact is determined primarily by the relatedness between the organism for which a model is being built and the organisms represented in the reconstruction data. These observations highlight principles that can help guide the selection of reconstruction data to improve metabolic network reconstruction.

## Results

### Source of reconstruction data affects model quality more than reconstruction method

It is reasonable to expect that using reconstruction data from a different source (e.g., BiGG instead of KEGG) would impact the models that are produced; however, it is not clear how large this impact is, especially relative to methodological changes (e.g., different gap-filling approaches). To establish the importance of reconstruction data in determining the composition of generated models, we selected two different reconstruction tools that have distinct methods and use data from different sources: Reconstructor (11), which uses data from KEGG and ModelSEED, and CarveMe (21), which uses data from BiGG. We also selected 10 different strains of bacteria that have manually curated models available in the BiGG repository (see Methods). We then generated two sets of models for the selected strains with each reconstruction tool, one with the tool’s default reconstruction data and the second with the other tool’s reconstruction data. In other words, we generated four total sets of models from all possible combinations of the two methods (Reconstructor or CarveMe) and two data sources (KEGG/ModelSEED or BiGG). We then compared the models based on which reactions were present or absent in each model (Fig 2A). The models were distinctly separated by the source of the reconstruction data, confirming that the source of data plays an important role in determining which reactions are included in the generated models. Additionally, for these two methods and data sources, there was greater separation between models by data source than by method, indicating that the source of data had a bigger impact than differences in method. These trends were also evident when we assessed how well the contents of the generated models recapitulated manually curated ones (Fig 2B). Overall, the source of reconstruction data substantially impacted what reactions were included in the generated models.

**Fig 2.**
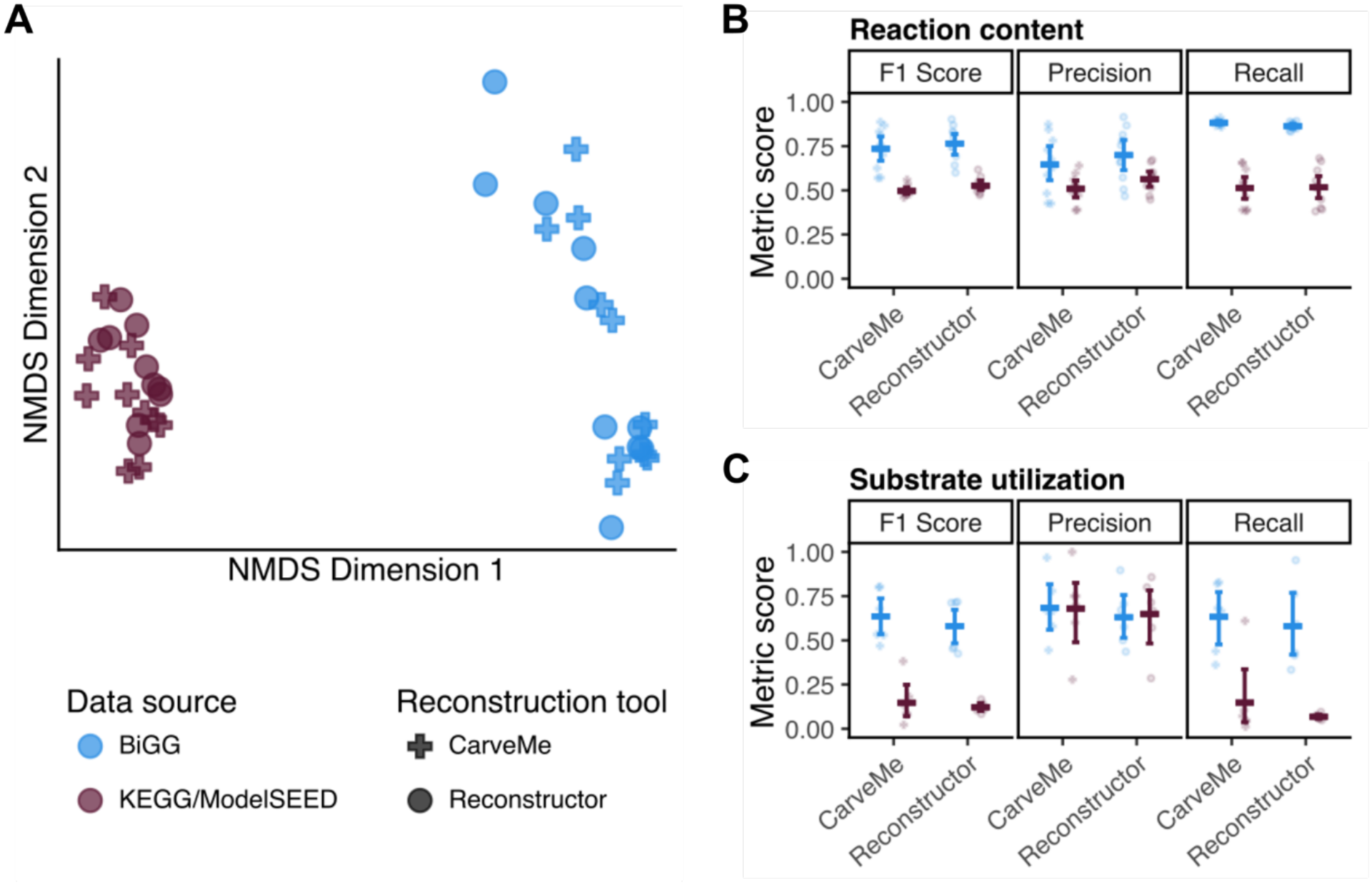
Impact of the source of reconstruction data on generated metabolic models. (A) Non-metric multidimensional scaling (NMDS) shows relative similarity between generated models based on the reactions they contain (normalized stress = 0.071). Models were generated using either Reconstructor or CarveMe and data from either BiGG or KEGG/ModelSEED. Data source refers to the source of data for both functional annotation and biochemical data. (B) Similarity between each generated model and a manually curated model for the same organism from BiGG. Similarity is based on the reactions contained in each model and is quantified using three metrics: F1 score, precision, and recall. (C) Similarity between substrate utilization predictions for each model and experimental data. For (B-C), bars indicate the mean and bootstrapped 95% confidence interval of each metric across the tested models.

After determining the impact of reconstruction data on model contents, we decided to assess if this impact extended to model predictions. For six of the bacteria strains which had available experimental data, we used the generated models to predict substrate utilization on various carbon sources. We then compared these predictions to the experimental data (Fig 2C). Similar to the contents of the models, we observed that the source of reconstruction data greatly affected the ability of the models to accurately predict carbon-source utilization, evident by the changes in F1 score. Interestingly, the effect was almost entirely on recall and not precision, which was a result of the models built with data from KEGG/ModelSEED predicting growth on far fewer substrates. Meanwhile, the difference in reconstruction methods had much smaller impact on the accuracy of the model predictions. Combined with the assessment of the model contents, these data demonstrate the importance of reconstruction data in the process of creating metabolic models.

### Functional annotation data differs between databases

The process of creating a genome-scale metabolic model involves two major components of reconstruction data: functional annotation data and biochemical data (Fig 1). One or both of these components may have contributed to the reconstruction differences we observed. Here, we focused our subsequent analyses on the functional annotation data and leave an in-depth exploration of the biochemical data to future work.

To determine why the source of data had such a large impact on the contents and predictions of generated models, we compared the functional annotation data between the different sources (Fig 3). Surprisingly, while KEGG had data for vastly more species of bacteria than BiGG (Fig 3A), BiGG captured more metabolic functions (Fig 3B). Additionally, in BiGG, each organism tended to have a higher number of genes that were linked to reactions than KEGG (Fig 3C), which helps explain why BiGG had greater coverage of metabolic functions despite containing fewer species. These results demonstrate that different sources of data vary in the scope, or amount, of data they contain. Consequently, one database may contain gene sequences or functions that the other does not. Even in areas where the databases overlap with one another, there are differences. For example, the gene *yqhD* in *Escherichia coli* was present in both KEGG and BiGG, but it was linked to a different set of reactions in each (Fig 3D). Together, these differences between databases suggest potential characteristics, such as the accuracy of annotations and the diversity of represented species, that likely influence the reconstruction process.

**Fig 3.**
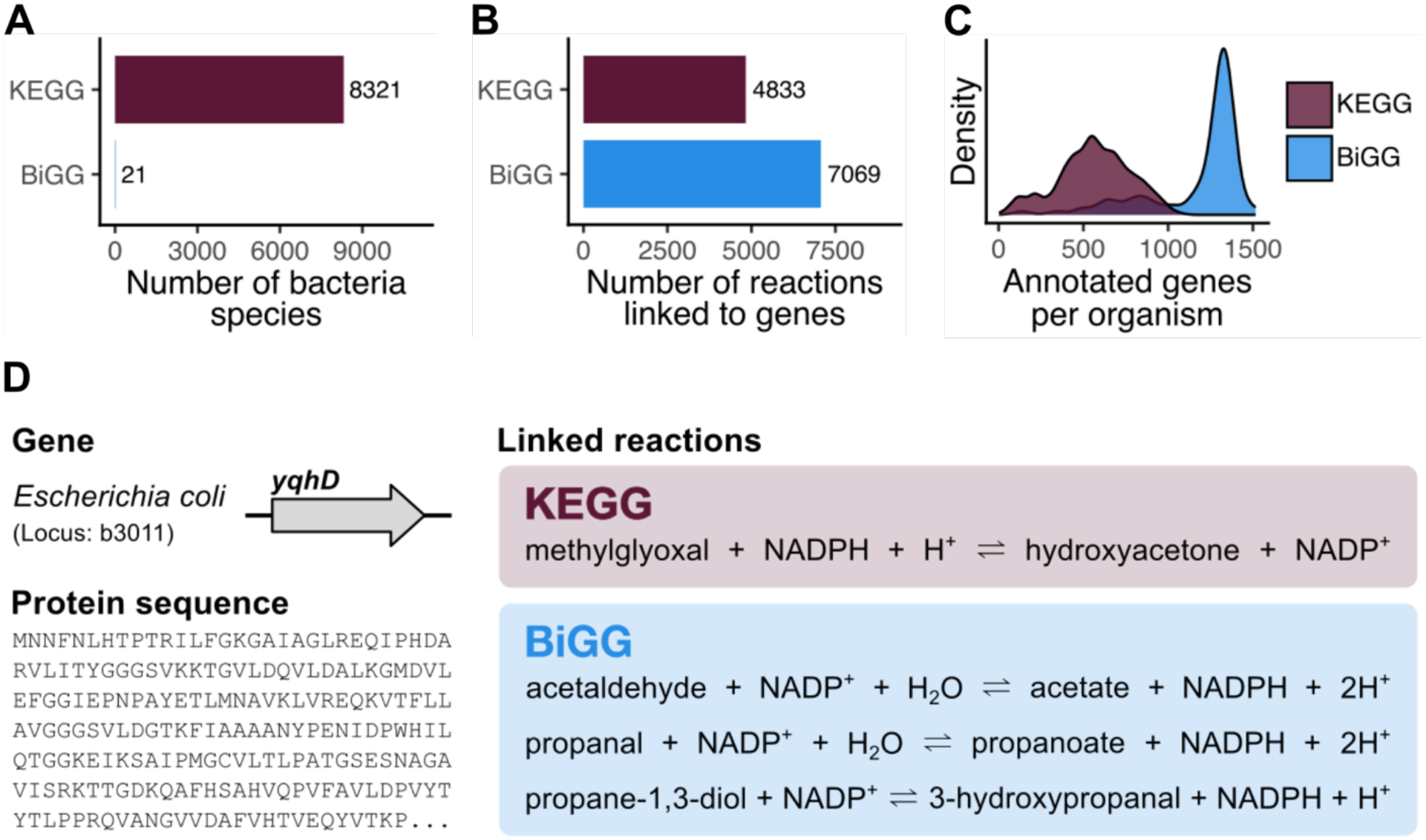
Differences in functional annotation data between KEGG and BiGG. (A) Total number of bacteria species with functional annotation data in each database. (B) Total number of unique reactions in each database that are linked to at least one bacterial gene. (C) Distribution of the number of annotated genes per organism in each database. Here, annotated genes are genes that are linked to at least one reaction in the database. (D) Example gene (*yqhD*) that is linked to different reactions in KEGG and BiGG.

### Automated reconstruction is sensitive to the accuracy of gene annotations

One potentially important parameter of functional annotation data is the accuracy of gene annotations (25). In other words, are the genes included in the data linked to the correct metabolic reactions? Some protein families have been shown to have a large number of incorrect annotations in public databases, even exceeding 80% misannotation in certain cases (33). Additionally, the frequency of misannotation are not consistent across databases (33). Consequently, we aimed to evaluate how differences in gene annotations affect automated network reconstruction. Starting with gene annotations from BiGG, we created several sets of annotations where each set had a fraction of genes that were randomly assigned a new annotation (i.e., each gene was randomly linked to a new set of reactions, see Fig 4A). Using Reconstructor and the same strains of bacteria as above, we generated models with each set of annotations. We compared the contents of the generated models to the corresponding manually curated models (Fig 4B-C). Unsurprisingly, we observed that introducing misannotations in the data reduced the similarity of the generated models to the curated ones. This effect was observed in all the species we tested and was remarkably linear, indicating that even small amounts of misannotation can reduce model quality. We also observed that on average, incorrect annotations reduced the accuracy of substrate utilization predictions (Fig 4D). These results demonstrate that the reconstruction process is sensitive to the accuracy of the functional annotation data it uses.

**Fig 4.**
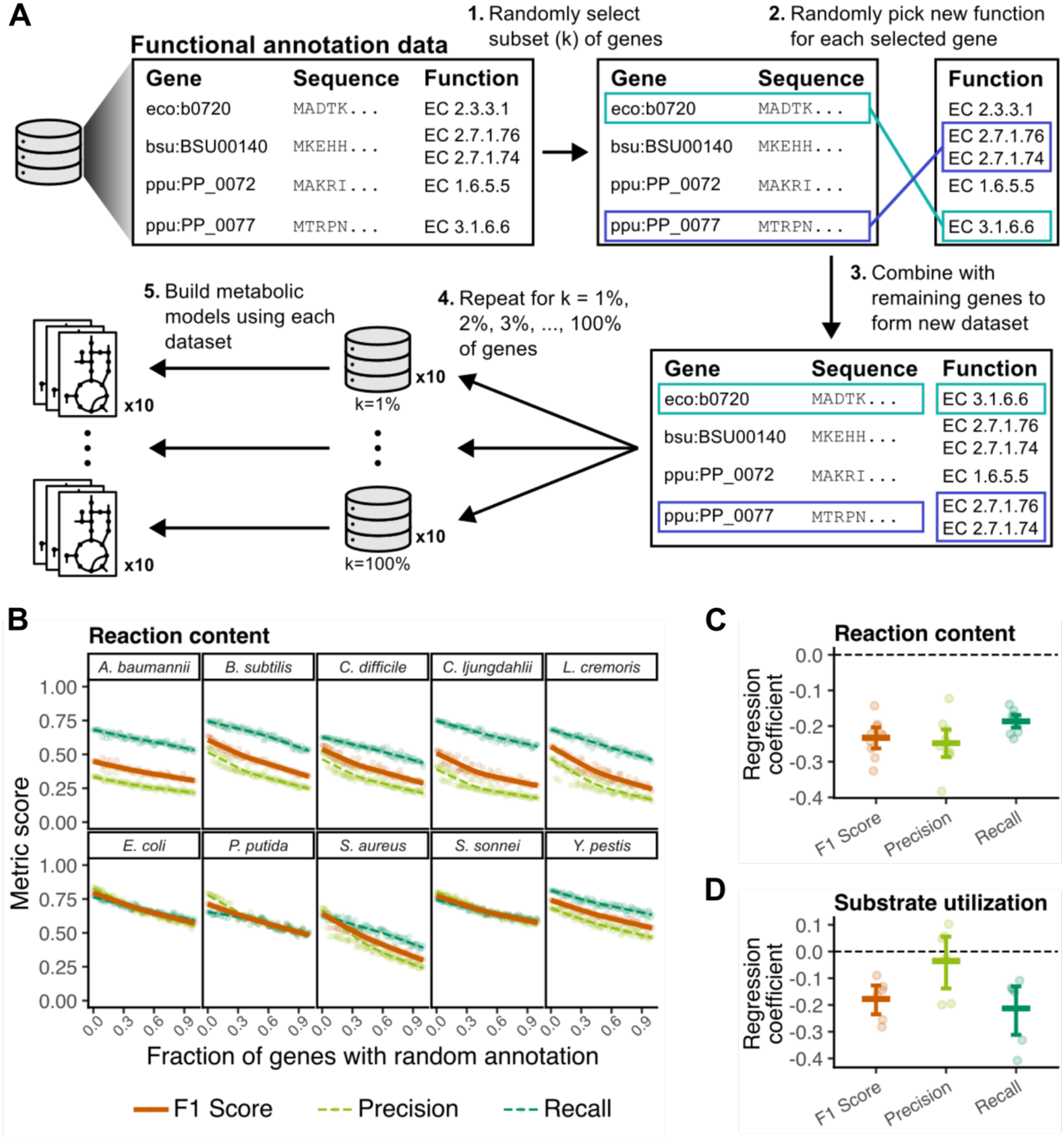
Effect of incorrect annotations in functional annotation data on generated metabolic models. (A) Approach to analyze the effect of incorrect annotations by randomly assigning new annotations to a subset of genes in the reconstruction data. (B) Similarity of generated models compared to manually curated ones as a function of the fraction of misannotated genes. Similarity is based on the reactions each model contains and is quantified using F1 score, precision, and recall. (C) Regression coefficients for each organism and metric from linear regression with the fraction of misannotated genes as the independent variable and the reaction content metric score as the dependent variable. Bars indicate the mean and 95% confidence interval of the coefficients from each organism. (D) Same as (C) but for substrate utilization predictions.

Given the observed sensitivity of the reconstruction process to incorrect annotations, we investigated whether a single annotation change in the functional annotation data was sufficient to impact the quality of a generated model. Employing a similar approach as before, we generated more than 1,000 models for each organism where each model was constructed using functional annotation data with a single annotation change (see Methods). We then compared the contents of these models to the corresponding manually curated ones (Fig 5A). While most single annotation changes had virtually no impact on the contents of the generated models, some changes affected the model contents considerably. A similar pattern was evident for the substrate utilization predictions of the generated models (Fig 5B). Of the misannotated genes that affected substrate utilization predictions, a handful had particularly strong effects when they were assigned a random annotation in the data (Fig 5C). For example, across three replicates, changing the annotation for the gene *fbp* from *Acinetobacter baumannii* (BiGG ID: ABAYE_RS05005) consistently led to *A. baumannii* models with exceptionally poor substrate utilization predictions (F1 score = 0.047 for all three models). Likewise, model predictions for *Pseudomonas putida* were sensitive to incorrect annotations for the gene *fbp* from *P. putida* (BiGG ID: PP_5040; F1 score = 0.19, 0.19, and 0.65 for three replicate models). Models for *Staphylococcus aureus*, on the other hand, were particularly sensitive to the *S. aureus* gene *gapA* (BiGG ID: USA300HOU_RS08940; F1 score = 0.27 for three replicate models). Interestingly, some annotation changes improved the substrate utilization predictions of generated models. For example, changing the annotation for either *pgk* (BiGG ID: BSU33930) or *gapA* (BiGG ID: BSU33940) from *Bacillus subtilis* resulted in *B. subtilis* models with better predictions overall (F1 score = 0.80 for three replicate models for each gene). These results demonstrate that a single incorrect gene annotation in reconstruction data can have a major impact on the models that are constructed.

**Fig 5.**
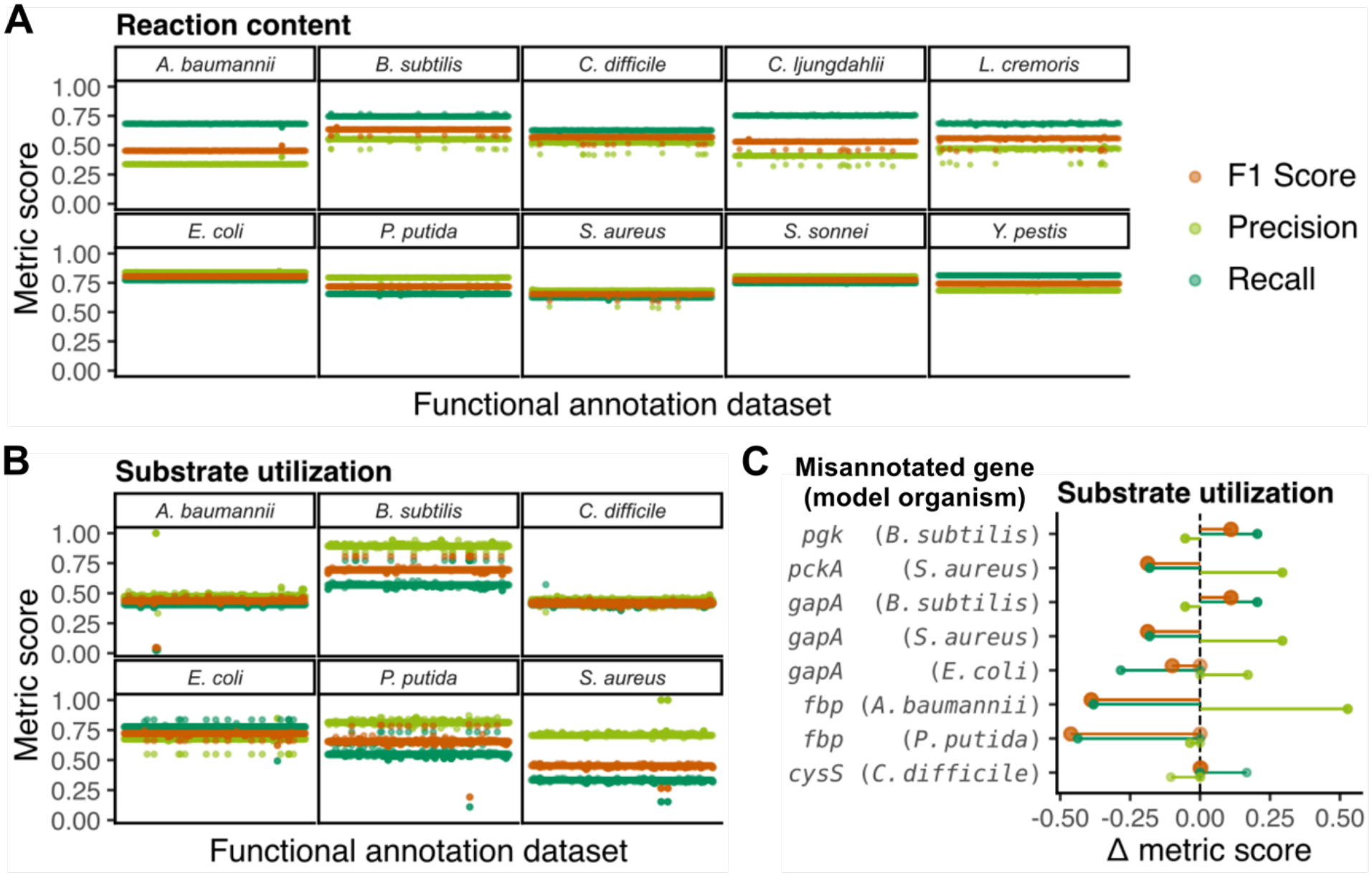
Effect of single annotation changes in functional annotation data on generated metabolic models. Models are evaluated based on the similarity of their (A) reaction contents to manually curated models and (B) substrate utilization predictions to experimental data. Each model was generated from functional annotation data with a single misannotated gene. Ordering of models along the x-axis is arbitrary. Similarity is quantified by F1 score, precision, and recall. (C) Genes that have the biggest effect on the quality of substrate utilization predictions for each model organism. The x-axis shows the difference in metric score from the baseline score (i.e., the score for the model built using data with no misannotations). There are three replicates for each gene.

### Data from closely related organisms improve network reconstruction quality

Another characteristic of functional annotation data is the diversity of organisms that the data include, or viewed another way, the taxonomic representation of the data. For example, some databases contain data (i.e., gene sequences and annotated functions) from a wide variety of organisms while others contain data from relatively few. Importantly, it has been noted that the completeness of computationally assigned annotations is influenced by taxonomy (34). This characteristic leads to the question, how much does the taxonomic representation of the reconstruction data matter for building metabolic network models? In particular, there are two aspects of taxonomic representation that we explored: 1) overall taxonomic coverage of the data and 2) taxonomic distance to the organism for which a model is being built (see Methods, Fig 6A). Taxonomic coverage reflects the number and diversity of the species that are represented in the reconstruction data. Taxonomic distance measures how closely related two organisms are to each other; in this case, the distance in question is between the organism for which a model is being built and the one nearest to it that is represented in the reconstruction data. Since closely related organisms have more similar genes and metabolic functions, we expect that using data from closely related organisms (i.e., low taxonomic distance) will produce better models. However, the magnitude of this impact is not known. Meanwhile, while higher taxonomic coverage may help recover more metabolic functions, it may also introduce pathways that are not present in the organism for which a model is being built. Thus, we sought to establish the relative importance of these two parameters (distance and coverage) and the direction of their effects.

**Fig 6.**
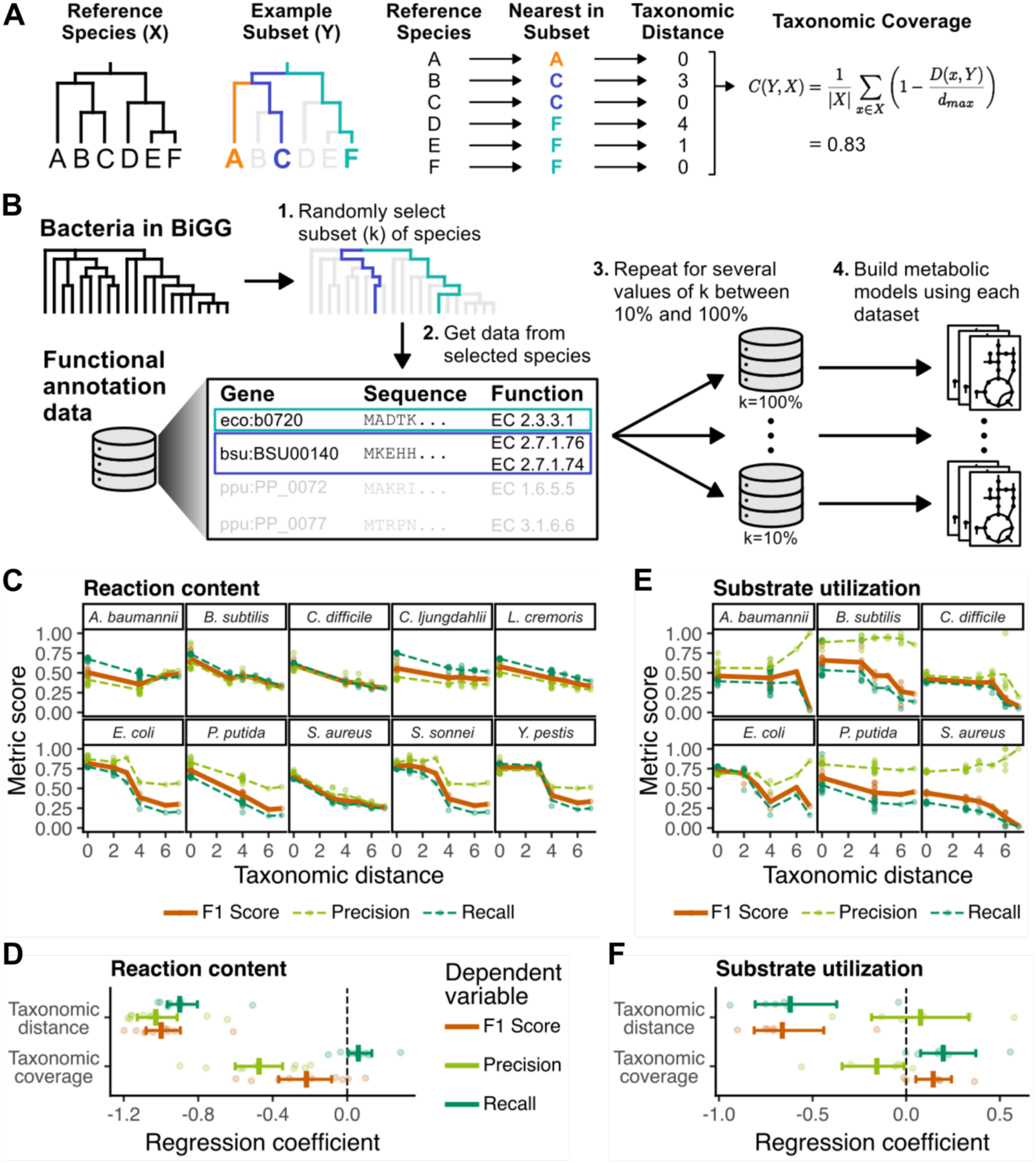
Importance of taxonomic parameters of the reconstruction data on automated network reconstruction. (A) Illustration of taxonomic distance and coverage for a hypothetical reference set of species and a selected subset of species. (B) Approach to measure the impact of taxonomic parameters of the reconstruction data by generating sets of data based on randomly selected subsets of bacteria. (C) Similarity between constructed models and manually curated ones as a function of taxonomic distance to the bacteria in the reconstruction data. Similarity is calculated based on the reactions included in each model. (D) Linear regression coefficients for taxonomic coverage and distance with the metrics from (C) as dependent variables. The regression coefficients indicate the effect of each taxonomic parameter on the reaction metrics. Bars indicate the mean and 95% confidence interval of the coefficients from each organism. (E) Similarity between substrate utilization predictions of the generated models and experimental data. (F) Same as (D) but using the metrics for substrate utilization prediction as dependent variables.

To measure the effects of taxonomic coverage and distance, we created different sets of reconstruction data with varying taxonomic representation (Fig 6B). Each set was created by pulling functional annotation data from a random subset of the bacterial species present in the BiGG database, ranging from 10 to 100% of the species in the database. We used the same biochemical data throughout to isolate the impact of the functional annotation data. We then generated models with Reconstructor using each of these sets of data and measured their similarity to manually curated models (Fig 6C). We saw a clear negative relationship between taxonomic distance and similarity to the curated models, which we quantified through linear regression (Fig 6D). As expected, taxonomic distance was negatively related to model quality, meaning that having data from a closely related organism (lower distance) leads to better quality (higher F1 score, precision, and recall). On the other hand, the effect of taxonomic coverage was much smaller. This effect was due to taxonomic coverage having simultaneous counter-balancing effects—greater coverage slightly improved recall but reduced precision. The combined effect (F1 score) on model contents for taxonomic coverage was negative. As before, we extended our analysis to substrate utilization predictions (Fig 6E-F). Taxonomic distance was again negatively correlated to overall model quality (F1 score). In this case, though, it only affected prediction recall and not precision. Meanwhile, taxonomic coverage again showed countering affects by increasing recall while decreasing precision. Surprisingly, the overall effect of increased coverage on model predictions was net positive, which was opposite the effect on model contents. This contrast indicates that there is not always a direct connection between model contents and predictions. All together, these results confirm the importance of using data from closely related organisms to produce higher-quality models. Additionally, they show that increased taxonomic coverage by itself does not clearly improve the contents of generated models but may improve their predictions.

## Discussion

Genome-scale metabolic models are valuable tools for understanding metabolism, but they are still limited by the difficulty of creating high-quality models. Automated tools for producing models have improved, but much of the focus has been on developing new methods rather than evaluating the data they use. Here, we demonstrated the impact that using different data can have on the contents and predictions of generated models. In the example we presented here between CarveMe and Reconstructor, the effect of different data was bigger than the effect of different methods. It should be noted that the relative sizes of these effects may not apply to all such comparisons depending on the degree of difference between methods and data sources. Additionally, it is possible that some methods may be more sensitive (or less sensitive) to changes in reconstruction data. Regardless, using different reconstruction data clearly impacts the reconstruction process.

Given the importance of reconstruction data, automated reconstruction tools should provide greater transparency and flexibility regarding the data that is used. As noted earlier, there is generally very little, if any, rationale or evidence provided for the use of data from one source over another. Additionally, it is not always clear what data is taken from what sources and how exactly that data is used by the tool. For most, if not all, currently existing reconstruction tools, the data is tightly integrated into the tool, giving users little flexibility to use different data that may be more suited to their objective. Even for the analyses in this paper, we had to modify both Reconstructor and CarveMe to be able to use them with different sets of data. Granted, the current lack of flexibility may be partly due to systematic differences between data sources, which make it difficult to design tools that can use data from multiple sources. However, here we showed that using different data can greatly affect network reconstruction, and thus, increased transparency and flexibility around reconstruction data is an area where all current reconstruction tools can improve.

The results we present highlight two important considerations for selecting reconstruction data—gene annotation quality and taxonomic representation—that can help guide future improvements to network reconstruction. When creating a metabolic network model for a particular organism, the reconstruction data should ideally include highly accurate annotation data for closely related organisms. Thus, one should consider both the accuracy of annotations within a database as well as the organisms that are included when selecting what data to use. There will likely be tradeoffs between annotation accuracy and taxonomic coverage, especially for organisms that are not well-studied. Such tradeoffs suggest that the reconstruction data may need to be chosen on a case-by-case basis depending on the identity of the organism for which a model is being built. This result further emphasizes the need to design tools so that users can choose what data they use. Additionally, functionality could even be included to help users identify what data is available and would be better suited for their objective.

While the results we present demonstrate the importance of reconstruction data, they are only a first step. In this work, we focused primarily on the functional annotation data used for network reconstruction and not the biochemical data. Previous work has shown that it is important to have consistent and complete biochemical data for gap-filling (35). However, other parameters remain to be explored for both the biochemical and functional annotation data. In particular, it would be valuable to evaluate the sensitivity of different annotation and gap-filling methods to changes in reconstruction data. For example, here we showed that Reconstructor could be sensitive to even one incorrect gene annotation, but other methods may show a different level of sensitivity to incorrect annotations. Further systematic assessments of the reconstruction process and its components is a promising direction to improve automated reconstruction tools and produce higher-quality models.

Reconstruction data, including both functional annotation data and biochemical data, are a critical component of the process to build metabolic network models. Since there are clear differences in the data from different databases, it is important to choose appropriate data when constructing a metabolic model. In fact, choosing what data to use may be even more important than choosing what method to use. Ideally, to build a model for a particular organism, one should have access to accurate functional annotation data from closely related organisms. Where such data are still missing, it emphasizes the value in continuing to functionally characterize genes from diverse species and compile those data in publicly available databases. As efforts continue to improve both reconstruction methods and the data they rely on, the quality of genome-scale metabolic network models and the predictions they yield will improve.

## Materials and Methods

### Selection of bacteria strains for analysis

Our objective was to assess how changes in reconstruction data impact the quality of metabolic network models across a range of bacterial taxa. To assess the quality of the models, we wanted to evaluate both their contents (i.e., what reactions are included) and their predictions (specifically, substrate utilization predictions) (14). In order to evaluate the quality of the model contents, we needed to select organisms with available, high-quality, curated models. Consequently, we selected bacterial species for which models are available from the BiGG repository, a collection of published models that have undergone significant manual curation and validation (26). We selected a total of ten phylogenetically diverse species with published models: *Acinetobacter baumanii* (36), *Bacillus subtilis* (37), *Clostridioides difficile* (38), *Clostridium ljungdahlii* (39), *Escherichia coli* (40), *Lactococcus cremoris* (41), *Pseudomonas putida* (42), *Shigella sonnei* (43), *Staphylococcus aureus* (44), and *Yersinia Pestis* (45). We ensured a balance between gram-positive and gram-negative bacteria by selecting five species of each. Additionally, we selected species that captured a range of distances between them and their closest-related species in BiGG. For example, the closest species to *Pseudomonas putida* is from the same class (Gammaproteobacteria) but not the same order (Pseudomonadales), while the closest to *Shigella sonnei* is from the same genus (*Shigella*). Of the ten species, we also selected six with available experimental substrate utilization data: *A. baumanii* (46), *B. subtilis* (37), *C. difficile* (47), *E. coli* (48), *P. putida* (*49*), and *S. aureus* (44).

### Local implementation of Reconstructor and CarveMe

The current versions of Reconstructor and CarveMe do not allow users to easily change the reconstruction data that they use. Thus, in order to analyze the impact of changing their reconstruction data, we made local copies of the source code for both Reconstructor (v1.2.0, commit ff8fb8c, https://github.com/csbl/reconstructor) and CarveMe (v1.6.6, commit 5b25ce5, https://github.com/cdanielmachado/carveme) from their GitHub pages. We made small changes to each tool to allow for the specification of different reconstruction data. Importantly, we did not change the core methodology of each tool, so our local implementations produce the same results as the released versions if given the same reconstruction data. The source code for our copies of these tools can be found on GitHub (https://github.com/abates20/bates-reconstruction-data) and have also been archived on Zenodo (https://doi.org/10.5281/zenodo.23701079).

### Evaluation of generated models

We evaluated the quality of generated models based on both the reactions they included and their predictions of substrate utilization (14). To assess the reactions included in a model, we compared its reactions to a high-quality, published model for the same strain of bacteria. We based this comparison on three metrics—recall, precision, and F1 score—which are based on the number of true positives (TP), false positives (FP), and false negatives (FN):

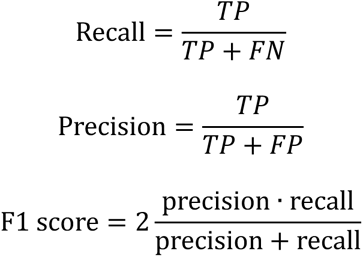

A reaction is considered a true positive if it is present in both the generated model and the published one. A reaction is a false positive if it is only present in the generated model. Finally, a reaction is a false negative if it is only present in the published model.

To assess the quality of substrate utilization predictions, we simulated growth of the models on each of the substrates included in the experimental data. For each simulation, we set the lower bound of the substrate exchange reaction to −10 mmol/g/h. For each other nutrient in the media, we set the lower bound of the corresponding exchange reaction to −1000 mmol/g/h (i.e., unlimited uptake). We considered growth to be positive on a particular substrate if the predicted biomass flux was >1 × 10^−6^. For the experimental data, we labeled growth vs no growth on a particular substrate using the same criteria as the paper where the data was originally reported. We then compared the predictions of the generated models to the experimental data using the same three metrics: recall, precision, and F1 score.

### Comparison of Reconstructor and CarveMe with different data sources

We took the reconstruction data from each tool (Reconstructor or CarveMe) and reformatted it to work with the other tool. In other words, we took the data from Reconstructor, which is derived from KEGG and ModelSEED, and reformatted it to work with CarveMe. We also took the data from CarveMe, which is derived from BiGG, and reformatted it to work with Reconstructor. We obtained an annotated genome (RefSeq annotations) for each of the ten selected bacterial species using the NCBI Datasets REST API (50). We then generated four models for each species—one model per combination of reconstruction tool (Reconstructor and CarveMe) and data source (KEGG/ModelSEED or BiGG). This process yielded 40 models total for this analysis.

To compare the contents of these models, we mapped all reaction IDs to the MetaNetX namespace using the MNXref 4.5 database (51). To ensure that differences between groups of models were not primarily driven by incomplete mapping between namespaces, we excluded any reactions from the MetaNetX namespace that could not be successfully mapped to both the ModelSEED and BiGG namespaces. Thus, the estimated differences between models that use different namespaces are a conservative estimate of the true difference in network structure between those models. We calculated the similarity of the generated models to each other and to the published models based on the presence or absence of reactions in each model. We calculated this similarity using the Jaccard distance and then visualized it using non-metric multidimensional scaling (NMDS). We also evaluated the quality of the generated models as described above.

### Analysis of impact of incorrect annotations

It is known that public sequence databases contain errors and biases (33,52,53), however the exact mechanism by which specific errors arise and the resulting distribution of annotation errors is hard to pinpoint. Thus, to model incorrect annotations, we took an unbiased approach and assumed that errors are essentially random. We modeled different levels of misannotation by assigning randomly selected functions to a subset of genes in the reconstruction data. In this case, a “function” refers to a set of reactions that are already linked to at least one gene in the data. Since using data from BiGG tended to yield higher quality models in our first analysis, we used data from BiGG for this analysis. We created 100 sets of reconstruction data with 1%, 2%, 3%,…, 100% of genes assigned random functions (Fig 4A). We then generated models using Reconstructor and each of these sets of reconstruction data. We also included a set of models generated with the normal (i.e., no random annotations) reconstruction data from BiGG. We evaluated the quality of these models as described above.

To assess the effect of single misannotations, we employed a similar approach where we assigned random functions to selected genes. This time, though, for each set of reconstruction data, we assigned a random function to only one gene. We created three sets of reconstruction data per gene in an effort to capture some of the variability that may arise depending on the random function that is assigned. Additionally, to reduce computational demand, we only constructed models using sets of reconstruction data where the misannotated gene came up as a hit in the initial BLAST step of the reconstruction process. If a gene does not show up as a hit in the initial BLAST step, then changing the gene’s annotation would have no effect on the reconstruction process and thus does not need to be included in the analysis. We evaluated the generated models as before by comparing their reaction contents to manually curated models and their substrate utilization predictions to experimental data.

### Assessment of effect of taxonomic representation

Similar to the analysis around annotation accuracy, we analyzed the effects of taxonomic representation by creating several different sets of reconstruction data. The data for each set was again derived from BiGG. For each set, we included functional annotation data (i.e., sequences and annotations) from only a fraction of the bacterial species in BiGG. We chose values ranging from 0.1 to 1.0 for the fraction of species, with 0.1 meaning that only 10% of species were included, and 1.0 meaning that all species were included. The biochemical data was consistent across all sets of reconstruction data to ensure successful gap-filling and isolate the effect of changing the functional annotation data. We then used each set of reconstruction data to build models and evaluated the quality of the models as described above. Additionally, we calculated the taxonomic coverage for each set of data and the taxonomic distance for each species of bacteria (see below). We estimated the effect of each of these two parameters using linear regression with the two parameters as independent variables and the quality metrics (recall, precision, and F1 score) as dependent variables.

### Calculation of taxonomic parameters

We investigated two taxonomic parameters of the reconstruction data: 1) taxonomic coverage and 2) taxonomic distance to a given organism (Fig 6A). Our method for quantifying these two parameters was inspired by previous work that incorporated taxonomic relatedness into measures of biodiversity (54). We denote taxonomic distance from a specific species, *x*, to a set of species, *Y*, as *D*(*x*, *Y*). It is the minimum path length along a taxonomic tree from *x* to any of the species in *Y*:

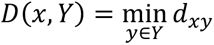

where *d_xy_* is the path length between species *x* and *y*. *d_xy_* ranges from 0 (i.e., *x* and *y* are the same species) to 8 (i.e., *x* and *y* are from different domains). Thus, *D*(*x*, *Y*) also ranges from 0 to 8. Given a set of species, *Y*, and a reference set of species, *X*, we can use taxonomic distance to calculate the coverage of *Y* with respect to *X*, which we denote *C*(*Y*, *X*):

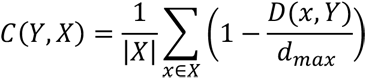

where *d_max_* is the maximum theoretical distance between any two species (which is 8 for species from different domains), and |*X*| is the number of species in the reference set. The minimum possible taxonomic coverage is 0, indicating that none of the species in *Y* are from the same domain as any of the species in *X*. The maximum possible coverage is 1, which means that every species in *X* is present in *Y*. In other words, taxonomic coverage reflects how well a set of species captures the taxonomic diversity in the reference set. In this work, we used all the bacterial species in BiGG as the reference set for calculating coverage.

